# Sexually divergent cortical control of affective-autonomic integration

**DOI:** 10.1101/2020.09.29.319210

**Authors:** Tyler Wallace, Derek Schaeuble, Sebastian A. Pace, Morgan K. Schackmuth, Shane T. Hentges, Adam J. Chicco, Brent Myers

**Author notes:** Equal contribution (Tyler Wallace, Derek Schaeuble). **Address for correspondence** Brent Myers, Ph.D., Department of Biomedical Sciences, Colorado State University, 1617 Campus Delivery, Fort Collins, CO 80523.

## Abstract

Depression and cardiovascular disease reduce quality of life and increase mortality risk. These conditions commonly co-occur with sex-based differences in incidence and severity. However, the biological mechanisms linking the disorders are poorly understood. In the current study, we hypothesized that the infralimbic (IL) prefrontal cortex integrates mood-related behaviors with the cardiovascular burden of chronic stress. In a rodent model, we utilized optogenetics during behavior and *in vivo* physiological monitoring to examine how the IL regulates affect, social motivation, neuroendocrine-autonomic stress reactivity, and the cardiac consequences of chronic stress. Our results indicate that IL glutamate neurons increase socio-motivational behaviors specifically in males. IL activation also reduced endocrine and cardiovascular stress responses in males, while increasing reactivity in females. Moreover, prior IL stimulation protected males from subsequent chronic stress-induced sympatho-vagal imbalance and cardiac hypertrophy. Our findings suggest that cortical regulation of behavior, physiological stress responses, and cardiovascular outcomes fundamentally differ between sexes.

## 1. Introduction

Major depressive disorder (MDD) and cardio-metabolic conditions including hypertension, glucose intolerance, and heart failure significantly contribute to global disease burden. Epidemiological evidence implicates life stressors as a risk factor for both MDD and cardiovascular disease (CVD) (Binder and Nemeroff, 2010; Grippo and Johnson, 2009; Myers et al., 2014b; Sgoifo et al., 2015). Furthermore, sex differences in the incidence of MDD, CVD, and MDD-CVD co-morbidity suggest that sex-specific factors contribute to outcomes (Goldstein et al., 2019). However, the biological basis for stress effects on health, particularly the integration of affective and physiological systems, is poorly understood. Human brain imaging studies indicate that ventral medial prefrontal cortex (vmPFC) activity associates with sadness and blood pressure reactivity, suggesting that top-down cortical control may integrate diverse aspects of mood and systemic physiology.

The vmPFC is involved in numerous cognitive and emotional processes (McKlveen et al., 2015; Myers-Schulz and Koenigs, 2012; Wood and Grafman, 2003). A subregion of the vmPFC, the subgenual cingulate cortex (BA25), is activated by sadness-provoking stimuli, responds to social isolation, and has reduced volume in MDD patients (Beckmann et al., 2009; Liotti et al., 2000; Vijayakumar et al., 2017). BA25 is also targeted for deep brain stimulation in patients with treatment-resistant depression, where larger volumes predict better treatment outcomes (Mayberg et al., 2005; Sankar et al., 2019). Although, broader investigation of BA25 activity in mood disorders has yielded mixed results with reports of both hyper- (Hamani et al., 2011; Mayberg et al., 2005) and hypo-activity (Drevets et al., 2008, 1997). Subgenual regions of vmPFC have also been identified as components of a central autonomic network monitoring visceral functions (Beissner et al., 2013; Gianaros and Sheu, 2009; Gianaros and Wager, 2015; Myers, 2016; Shoemaker et al., 2016). Furthermore, recent pharmacological studies employing glutamate uptake inhibitors in non-human primates have implicated BA25 activity in reduced reward motivation (Alexander et al., 2019) and enhanced threat-related autonomic responses (Alexander et al., 2020). The rodent putative anatomical homolog of BA25, the infralimbic cortex (IL) (Öngür et al., 2003; Roberts and Clarke, 2019; Uylings et al., 2003; Vertes, 2004), innervates limbic and stress-regulatory nuclei including the amygdala, thalamus, and hypothalamus (Gabbott et al., 2005; Myers et al., 2016, 2014a; Wood et al., 2018). Additionally, IL stimulation reduces passive coping and increases pyramidal neuron spine density in male rodents (Fuchikami et al., 2015). Moreover, knockdown of IL glutamatergic output exacerbates chronic stress effects on hypothalamic-pituitary-adrenal (HPA) axis reactivity and vascular function (Myers et al., 2017; Schaeuble et al., 2019). However, the potential sex-specific roles of IL activity to integrate socio-motivational behaviors with physiological stress reactivity and the cardiac outcomes of chronic stress remain to be determined.

To identify how the male and female vmPFC coordinates mood-related behaviors and cardiovascular outcomes, genetically-identified vmPFC projection neurons received temporally-specific stimulation in an *in vivo* rodent model. Specifically, channelrhodopsin-2 (ChR2) was expressed under the calcium/calmodulin-dependent protein kinase type II α (CaMKIIα) promoter in the IL to permit optogenetic activation of IL pyramidal neurons in both male and female rats (Wood et al., 2018). This approach was combined with measures of place preference and social behavior to examine affective valence and sociability. Behavioral assessment was followed by measures of physiological stress reactivity, including radiotelemetry and echocardiography over the course of chronic variable stress (CVS). Ultimately, these findings identify the IL as an affective-autonomic integrator that links motivation and stress responding divergently in males and females.

## 2. Methods

### 2.1 Animals

Age-matched adult male and female Sprague-Dawley rats were obtained from Envigo (Denver, CO) with male rat weight ranging from 250-300 g and female from 150-200 g. After stereotaxic surgery, rats were housed individually in shoebox cages with cardboard tubes for enrichment in a temperature- and humidity-controlled room with a 12-hour light-dark cycle (lights on at 07:00h, off at 19:00h) and food and water *ad libitum*. In accordance with ARRIVE guidelines, all treatments were randomized and experimenters blinded. All procedures and protocols were approved by the Colorado State University Institutional Animal Care and Use Committee (protocol: 16-6871A) and complied with the National Institutes of Health Guidelines for the Care and Use of Laboratory Animals. Signs of poor health and/or weight loss ≥ 20% of pre-surgical weight were *a priori* exclusion criteria. These criteria were not met by any animals in the current experiments; however, animals were removed from experimentation if fiber optic or radiotelemetry devices failed.

### 2.2 Microinjections

Rats were anesthetized with isoflurane (1-5%) and administered analgesic (0.6 mg/kg buprenorphine-SR, subcutaneous). Rats received bilateral microinjections (Males 1.5 - 2 µL, Females 0.75 – 1.25 µL) of adeno-associated virus (AAV) into the IL (males: 2.7 mm anterior to bregma, 0.6 mm lateral to midline, and 4.2 mm ventral from dura, females: 2.3 mm anterior to bregma, 0.5 mm lateral to midline, and 4 mm ventral from dura). These volumes correspond with prior studies utilizing viral vector transduction in rat vmPFC (Ferenczi et al., 2016; Ji and Neugebauer, 2012; Wood et al., 2018). AAV5-packaged constructs (University of North Carolina Vector Core, Chapel Hill, NC) either expressed yellow fluorescent protein (YFP) or ChR2 conjugated to YFP under the CaMKIIα promoter to achieve pyramidal cell-predominant expression (Wood et al., 2018). All microinjections were carried out with a 25-gauge, 2-µL microsyringe (Hamilton, Reno, NV) using a microinjection unit (Kopf, Tujunga, CA) at a rate of 5 minutes/µL. The needle was left in place for 5 minutes before and after injections to reduce tissue damage and allow diffusion. Skin was closed with wound clips that were removed 2 weeks after injections and animals were allowed at least 6 weeks for recovery and ChR2 expression.

### 2.3 Electrophysiology

Adult male rats (n = 8) were injected with AAV constructs as described above and, after 8-12 weeks, exposed to 5% isoflurane prior to decapitation and brain removal. As previously described (Rau and Hentges, 2017), brains and sections were collected in ice-cold artificial CSF (aCSF) consisting of the following (in mM): 126 NaCl, 2.5 KCl, 1.2 MgCl_2_ Ʌ 6H_2_O, 2.4 CaCl_2_ Ʌ 2H_2_O, 1.2 NaH_2_PO_4_, 11.1 glucose, and 21.4 NaHCO_3_, bubbled with 95% O_2_ and 5% CO_2_. Coronal slices containing the IL were cut at a thickness of 240 μm using a model VT1200S vibratome (Leica Microsystems, Buffalo Grove, IL). After resting 1 hr at 37°C in aCSF, slices were transferred to the recording chamber and perfused with oxygenated 37° C aCSF at a 2 ml/min flow rate. For whole-cell recordings, the internal recording solution contained the following (in mM): KCL 57.5, K-methyl sulfate 57.5, NaCl 20, MgCl_2_ 1.5, HEPES 5; EGTA 0.1; ATP 2; GTP 0.5, and phosphocreatine 10. The pH was adjusted to 7.3. Recording electrodes had a resistance of 2 – 4 MΩ when filled with this solution. IL pyramidal neurons were identified for recording based on the expression of ChR2-YFP under the control of CaMKIIα. Whole-cell patch-clamp recordings were acquired in voltage-clamp at a holding potential of −60 mV using an Axopatch 200B Amplifier (Molecular Devices, San Jose, CA). Current-clamp recordings were acquired while holding current at 0 pA. Electrophysiological data were collected and analyzed using Axograph X software on a Mac OS X operating system (Apple, Cupertino, CA). Light activation of IL neurons expressing ChR2 occurred via 473 nm LED (Thorlabs, Newton, NJ) 1.1 mW light pulse driven by a LEDD1B driver (Thorlabs, Newton, NJ) triggered through the TTL output on an ITC-18 computer interface board (HEKA Instruments, Holliston, MA). Current-clamp experiments utilized 5, 10, and 20 Hz stimulation frequencies for 5 min bouts. Recordings were excluded if access resistance exceeded 10 Ω during recording.

### 2.4 Radiotelemetry implantation

A subset of rats was instrumented with ECG-enabled radiotelemetry transmitters (HD-S11 F0, Data Sciences International, St. Paul, MN) as previously described (Flak et al., 2011; Goodson et al., 2017; Schaeuble et al., 2019). Briefly, rats were anesthetized with inhaled isoflurane anesthesia (1-5%) and given a subcutaneous injection of analgesic (0.6 mg/kg Buprenorphine-SR) and an intramuscular injection of antibiotic (5 mg/kg gentamicin). The descending aorta was exposed via an abdominal incision, allowing implantation of a catheter extending from the transmitter. The catheter was secured with tissue adhesive (Vetbond; 3M Animal Care Products, St. Paul, MN) applied over a cellulose patch. ECG leads were passed through abdominal musculature and sutured subcutaneously above the rib cage and pectoral muscles. The transmitter body was then sutured to the abdominal musculature, followed by closure of the abdominal and skin with suture and wound clips, respectively. Rats then recovered for 2 weeks before wound clips were removed.

### 2.5 Fiber optic cannulas

Rats were anesthetized with isoflurane (1-5%) followed by analgesic (0.6 mg/kg buprenorphine-SR, subcutaneous) and antibiotic (5 mg/kg gentamicin, intramuscular) administration. Bilateral fiber-optic cannulas (flat tip 400/430 μm, NA = 0.66, 1.1 mm pitch with 4.5 mm protrusion for males and 4.2 mm protrusion for females; Doric Lenses, Québec, Canada) were aligned with the IL injection sites and lowered to the ventral PL/dorsal IL approximately 1 mm dorsal to the injection to enable optic stimulation of the IL. Cannulas were secured to the skull with metal screws (Plastics One) and dental cement (Stoelting, Wood Dale, IL). Skin was sutured and, following 1 week of recovery, rats were handled daily and acclimated to the stimulation procedure for another week before experiments began. Rat handling and cannula habituation continued daily throughout experiments.

### 2.6 Optogenetic stimulation

Light pulses (1.0 - 1.3 mW, 5 ms pulses, 10 or 20 Hz) were delivered through a fiber-optic patch cord (240 μm core diameter, NA = 0.63; Doric Lenses) connected to a 473 nm LED driver (Doric Lenses). Optic power was measured with a photodiode sensor (PM160, Thorlabs Inc, Newton, NJ) at the cannula fiber tip. Initial male experiments used 20 Hz stimulation but, following the results of slice electrophysiology experiments, all subsequent studies used 10 Hz stimulation. Rats received a single 20-minute session of stimulation (1 min on/1 min off) in their homecage the week prior to stress exposure. In total, rats received self-determined stimulation in the stimulation zone of RTPP day 2, throughout the social behavior assay, and during acute stress exposure (restraint and novel environment), all prior to CVS. Immediately before and after CVS, ultrasound measures were conducted with optics both off and on to isolate long-term structural changes and acute functional changes. Further optic details are elaborated in the specified sections below.

### 2.7 Estrous cycle cytology

All female rats went through experiment 3 simultaneously, housed in the same room and randomly cycling. Immediately following acute measures of behavior and physiology, vaginal cytology was examined to approximate the estrous cycle stage. A damp (deionized water) cotton swab was used to collect cells from the vaginal canal and place them onto a glass slide. When dried, slides were viewed under a 10x objective light microscope by a minimum of two blind observers and were categorized as proestrus, estrus, metestrus, or diestrus (Cora et al., 2015; Smith et al.; Solomon et al., 2015).

### 2.8 Real-time place preference

The real-time place preference (RTPP) assay was used to assess the valence of IL stimulation (Bimpisidis et al., 2020; Stamatakis and Stuber, 2012). Cannulas were connected via patch cords to LEDs for light delivery and rats placed in a custom-made fiberglass arena with two chambers connected by a corridor (chambers: 15 x 15’’, corridor: 8 x 6’’, 15’’ deep). Rats explored the arena for 10 minutes on two consecutive days. The first day was a habituation day and no stimulation was delivered on either side. On the second day, rats received LED-generated 470 nm light pulses upon entry and throughout the time spent in the assigned stimulation side. Stimulation stopped when rats exited the assigned stimulation side but re-commenced upon re-entry. Thus, rats determined the amount of stimulation received through time spent in the stimulation side. Trials were recorded by a camera mounted above the arena and animal position was tracked by Ethovision software (Noldus Information Technologies) for automated optic hardware control. Stimulation side assignment was counterbalanced and animal testing was randomized. The time rats spent in the stimulation side was divided by the total time and multiplied by 100 to generate a percentage of time spent in the stimulation side.

### 2.9 Social behavior

A modified version of the 3-chambered social behavior assay was used to accommodate optic patch cords (Felix-Ortiz and Tye, 2014; Moy et al., 2004). To examine social interaction, each rat was connected to a patch cord and placed in a black rectangular fiberglass arena (36 x 23’’, 15.8’’ deep). Initially, the arena was empty and experimental rats were allowed to explore for 5 minutes without optic stimulation. The experimental rat was then returned to their home cage while an empty enclosure (ventilated with small round openings) was placed on one side of the arena, defined as the object, and an identical enclosure containing an age- and sex-matched conspecific was placed on the other side of the arena, defined as the social cage. The experimental rat was then placed in the middle of the arena and allowed to explore freely for 10 minutes with 5 ms pulsatile stimulation delivered throughout to quantify social motivation. The experimental rat was then placed again into its homecage while the empty enclosure was replaced with a new enclosure containing a novel age- and sex-matched conspecific. The experimental rat freely explored for 10 minutes while receiving optic stimulation to assess social novelty preference. Behavior was recorded with an overhead camera and interactions were defined as nose pokes onto cages and scored by a treatment-blinded observer. The duration of interactions was divided by the total time of each interaction period and multiplied by 100 to give a percent interaction value. Sides for object cage, social cage, and novel cage were counterbalanced and animal order randomized.

### 2.10 Novel environment

In rats with radiotelemetry implants, baseline cardiovascular measurements were collected the weekend prior to novel environment stress. Rats were then exposed to a 30-minute novel environment stressor to assess acute cardiovascular reactivity. During novel environment, rats were connected to fiber-optic patch cords for stimulation and placed into a brightly-lit semi-transparent arena. Radiotelemetry receiver pads (Data Sciences International, St. Paul, MN) were arrayed under the arena with 10 Hz optic stimulation during the stressor to record hemodynamics and activity in 1-minute bins. Heart rate (HR), mean arterial pressure (MAP), systolic arterial pressure (SAP), and diastolic arterial pressure (DAP) were collected and analyzed with Ponemah software (Version:6.4x Data Sciences International).

### 2.11 Restraint stress

Restraint was used to examine neuroendocrine responses to acute stress. Rats were placed in plastic film decapicones (Braintree Scientific, Braintree, MA) and connected to fiber-optic patch cords for optic stimulation throughout the 30-minute restraint. Blood samples (approximately 250 μL) were collected by tail clip at the initiation of restraint with additional samples taken 15 and 30 min after (Vahl et al., 2005). At the conclusion of restraint, patch cords were disconnected and rats returned to their homecage with recovery blood samples collected at 60 and 90 min after the initiation of restraint. Blood glucose was determined with Contour Next EZ glucometers (Bayer, Parsippany, NJ) and 2 independent readings for each time point were averaged. Blood samples were centrifuged at 3000 X g for 15 minutes at 4° C and plasma was stored at –20° C until radioimmunoassay (RIA). Plasma corticosterone levels were measured with an ^125^I RIA kit (MP Biomedicals, Orangeburg, NY) as previously described (Myers et al., 2017). All samples for each sex were run in duplicate and all time points were run in the same assay. RIA intra-assay coefficient of variation was 8.6% and interassay 13.6%.

### 2.12 Chronic variable stress

Chronic variable stress was comprised of twice daily (AM and PM) repeated and unpredictable stressors presented in a randomized manner including: exposure to a brightly-lit open field (28.5 x 18’’ 13’’ deep, 1 hour), cold room (4° C, 1 hour), cage tilt (45°, 1 hour), forced swim (23° to 27° C, 10 minutes), predator odor (fox or coyote urine, 1 hour), restraint (1 hour), and shaker stress (100 rpm, 1 hour). Additionally, overnight stressors were variably included, comprised of damp bedding (400 mL water) and overnight light. During the 2 weeks of CVS, rats were weighed every 3-4 days to monitor body weight.

### 2.13 Spectral analysis of heart rate variability

Throughout CVS, resting homecage recordings of heart rate variability (HRV) were collected and analyzed using Ponemah software (Version:6.4x Data Sciences International, St. Paul, MN). Using guidelines for heart rate variance (Electrophysiology, 1996) in the frequency domain, low (LF) and high frequency (HF) components were collected during two-hour periods in the AM and PM and averaged within treatment groups. Recordings were sampled at least one hour after stressors and did not follow overnight stressors, swim, or restraint. Spectral analysis is used to measure autonomic balance as LF predominantly represents sympathetic and HF predominantly parasympathetic contributions to HRV (Electrophysiology, 1996; Sgoifo et al., 2015). Accordingly, LF/HF represents net cardiac sympathetic drive.

### 2.14 Echocardiography

Left ventricle structure and function were assessed before and after CVS via echocardiography. In preparation, rats were anesthetized with inhaled isoflurane (5% induction, 2% maintenance), connected to fiber optic patch cord, and shaved over the ventral thorax. A 12-mHz pediatric transducer connected to a Phillips XD11 ultrasound system was used to image the heart in transverse (parasternal short axis) and 4-chamber angles. M-mode echocardiograms were utilized to measure left ventricular end-systolic and end-diastolic chamber dimensions and anterior/posterior wall thickness (Chicco et al., 2008; Le et al., 2014). Once the measurements were taken for a rat, optics were turned on (1.3 mW, 5 ms pulses, 10 Hz) and a second exam was conducted to investigate acute stimulation-induced changes. Total exam time for each trial was approximately 5 minutes with approximately 2 minutes of optic stimulation.

### 2.15 Tissue collection

At the conclusion of experiments, rats were given an overdose of sodium pentobarbital and perfused transcardially with 0.9% saline followed by 4.0% paraformaldehyde in 0.1 M PBS. Brains were removed and post-fixed in 4.0% paraformaldehyde for 24 h at room temperature, followed by storage in 30% sucrose in PBS at 4 °C. Coronal sections were made on a freezing microtome at 30 μm thickness and then stored in cryoprotectant solution at −20 °C until processing. A subset of rats received optogenetic stimulation prior to tissue collection. Rats were tethered to a fiber optic patch cord and received 5 minutes of optic stimulation (1 mW, 5 ms pulses, 10 Hz) followed by 90 minutes of recovery for immediate-early gene (c-Fos) expression prior to euthanasia, as described above.

### 2.16 Immunohistochemistry and microscopy

For fluorescent labeling of c-Fos, coronal brain sections were removed from cryoprotectant and rinsed in PBS (5 x 5 min) at room temperature. Sections were then placed in blocking solution (PBS, 0.1% bovine serum albumin, and 0.2% Triton X-100) for 1 hour. Next, sections were incubated overnight in rabbit anti-c-Fos primary antibody (1:200 in blocking solution, Cell Signaling Technologies, Ab #2250). Following overnight incubation in primary antibody, sections were rinsed in PBS (5 x 5 min) and incubated in donkey anti-rabbit Cy5 secondary (1:1000 in PBS, Jackson ImmunoResearch, AB_2340607) for 1 hour. The tissue was then washed (5 x 5 min in PBS), mounted with polyvinyl medium, and cover slipped for imaging. For chromogen labeling of c-Fos positive nuclei, sections were first rinsed in PBS (5 x 5 min) at room temperature. Sections were then incubated in 1% hydrogen peroxide in PBS for 10 minutes followed by a second PBS (5 x 5 min) wash. Sections were then placed in blocking solution for 1 hour. Following blocking, sections were incubated overnight in rabbit anti-c-Fos primary antibody (1:2000 in blocking solution, Abcam, ab190289). Sections were then rinsed in PBS (5 x 5 min) followed by a 1 hour incubation in biotinylated goat anti-rabbit secondary (1:500 in PBS, Vector Laboratories, BA-1000). Sections were rinsed in PBS (5 x 5 min) followed by a 1 hour incubation in Vectastain ABC Solution (1:1,000; Vector Laboratories). Sections were rinsed again prior to incubation in diaminobenzidine and hydrogen peroxide (0.02% diaminobenzidine/0.09% hydrogen peroxide in KPBS) for 10 min. Following incubation, slices were rinsed, slide mounted, dehydrated in graded ethanol, and cover slipped.

To determine injection placement, YFP was imaged with a Zeiss Axio Imager Z2 microscope using the 10x objective, while YFP and c-Fos dual fluorescence were acquired as tiled 20x objective images. For quantification of c-Fos positive cells, slides were imaged in brightfield with the 10x objective.

### 2.17 Quantification of c-Fos

To quantify the number of c-Fos positive cells, 3-4 IL micrographs adjacent to visible cannula tracts were imaged and positive cells within a half-width 10x frame were hand-counted by a treatment-blind observer.

### 2.18 Experimental design

Timelines for experiments are outlined in figure 1. Experiment 1 was comprised of 2 cohorts of male rats (n = 13-14 each) to yield 10 YFP and 17 ChR2. For this experiment, real-time place preference and social behavior were assessed prior to restraint stress with 20 Hz used for all stimulation. A parallel group of injected male rats (n = 8) were used for slice electrophysiology. Experiment 2 consisted of 3 cohorts of male rats (n = 7-8 each) for a total of 12 YFP and 10 ChR2. A subset of these rats was equipped with radiotelemetry transmitters (YFP n = 7, ChR2 n = 8) and all underwent real-time place preference and social behavior prior to novel environment with 10 Hz stimulation. Afterward, rats in experiment 2 underwent echocardiography before and after CVS. Experiment 3 was designed as a single cohort of female rats (n = 29) with 12 YFP and 17 ChR2. A subset of these rats was equipped with radiotelemeters (YFP n = 7, ChR2 n = 8). Real-time place preference and social motivation were followed by novel environment and restraint with 10 Hz stimulation for all measures. Female rats also underwent echocardiography before and after CVS. A subset of rats in experiments 2 and 3 (n = 6-7/group/sex) received optic stimulation prior to tissue collection to verify neuronal activation.

**Figure 1:**
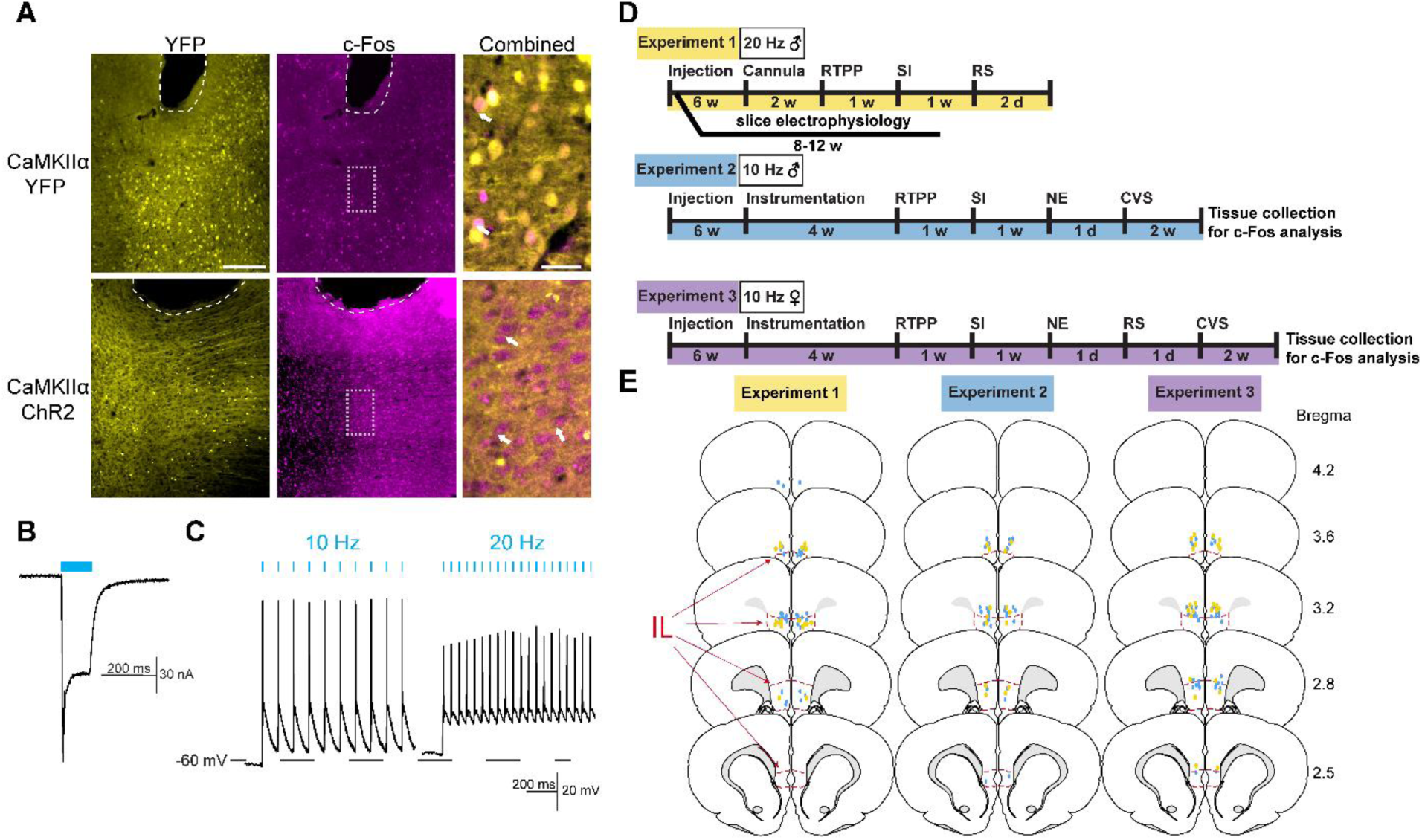
Approach validation and experimental design. (**A**) Injection of AAV-packaged constructs led to expression of cytosolic YFP or membrane-targeted ChR2-YFP under the CaMKIIα promoter. Blue light stimulation in ChR2 females (1 mW, 10 Hz, 5 ms pulses for 5 min) led to robust expression of the immediate-early gene marker c-Fos. Bregma +2.8mm. White arrows indicate representative c-Fos-positive nuclei. Dashed white lines indicate fiber tip. Scale bar: 200 µm and 40 µm for combined. (**B**) Voltage-clamp recordings from male IL-containing slices illustrated light-evoked (1.1 mW, 100 ms pulse) depolarizing current. (**C**) Current-clamp recordings found stimulation-locked spiking with 10 and 20 Hz stimulation (1.1 mW, 5 ms pulse). (**D**) Experimental timelines. RTPP: real-time place preference, SI: social interaction, RS: restraint stress, NE: novel environment, CVS: chronic variable stress. (**E**) Optic fiber placements (YFP: yellow, ChR2: blue) were mapped within or immediately dorsal to the IL (red outline). Coronal sections adapted from Swanson Rat Brain Atlas (3rd edition).

### 2.19 Data analysis

Data are expressed as mean ± standard error of the mean. Data were analyzed using Prism 8 (GraphPad, San Diego, CA), with statistical significance set at p < 0.05 for rejection of null hypotheses. Stimulation induced c-Fos was analyzed with Welch’s unpaired t-test comparing virus groups. RTPP stimulation preference was assessed via repeated measure two-way analysis of variance (ANOVA) with virus and day (repeated) as factors. In the case of significant main or interaction effects, Sidak’s multiple comparisons were run to determine group differences. Social motivation and novelty preference were assessed with unpaired t-tests to compare virus groups within social, object, novel rat, or familiar rat interactions. Total distance traveled during RTPP and social interaction was assessed with unpaired t-tests comparing virus groups. Stress responses over time (corticosterone, glucose, HR, MAP, SAP, DAP, and HRV spectra) were analyzed using mixed-effects analysis with virus and time (repeated) as factors, followed by Fisher’s post-hoc if significant main or interaction effects were present. Baseline HR and MAP, as well as mean activity during novel environment, were assessed with Mann-Whitney U non-parametric test comparing virus groups. Fractional shortening and ventricular structure were assessed with paired Wilcoxon signed-rank tests comparing pre- and post-CVS or optic status within virus groups.

## 3. Results

### 3.1 Validation and design

AAV viral vectors were targeted to the IL for expression of membrane-targeted ChR2-YFP or cytosolic YFP under the pyramidal neuron promoter, CaMKIIα. Fiber optic cannulas implanted in the IL permitted selective stimulation of IL glutamatergic neurons (**Fig. S1**), as previously reported (Fuchikami et al., 2015; Wood et al., 2018). In female rats, representative c-Fos labeling demonstrated *in vivo* neural activation in response to light-stimulation (1 mW, 10 Hz, 5 ms, 5 min) (**Fig. 1A**). c-Fos quantification in both male and female rats (**Fig. S2**) further validated stimulation efficacy. Whole-cell patch-clamp recordings in slice demonstrated light-evoked depolarizing current in male IL neurons expressing ChR2 (**Fig. 1B**). Further, 10 Hz stimulation led to high-fidelity action potential generation. 20-Hz light pulses induced action potential firing, but also increased resting membrane potential (**Fig. 1C**). These results, combined with our previous studies quantifying increased immediate-early gene expression after 20 Hz stimulation (Wood et al., 2018), suggest that both 10 Hz and 20 Hz stimulation activate IL CaMKIIα-positive neurons. Although, 10 Hz stimulation is more in line with the reported 4-10 Hz intrinsic firing rate of IL pyramidal neurons (Homayoun and Moghaddam, 2007; Ji and Neugebauer, 2012). The experimental design (**Fig. 1D**) is detailed in the methods. Male and female rats received optogenetic stimulation during behavioral and physiological measures in separate experiments. At the conclusion of all experiments, the placement of viral injections and fiber optics was determined. Only animals with cannula placement verified to be within or immediately (within 0.5 mm) dorsal to the IL were included in analyses (**Fig. 1E**).

### 3.2 Real-time place preference

RTPP was used to examine the affective valence of stimulating IL glutamatergic neurons. Group mean heat maps illustrate time spent in the stimulation versus non-stimulation chambers (**Fig. 2A**). Male rats showed a preference for the stimulation chamber (**Fig. 2B**) with 20 Hz (repeated-measures 2-way ANOVA: stimulation x ChR2 F_(1, 23)_ = 15.04, *p <* 0.001) and 10 Hz stimulation (repeated-measures 2-way ANOVA: stimulation F_(1, 20)_ = 6.03, *p* < 0.05, ChR2 F_(1, 20)_ = 9.04, *p <* 0.01, stimulation x ChR2 F_(1, 20)_ = 6.43, *p <* 0.05). Specifically, Sidak’s *post hoc* indicated a preference for the 20 Hz and 10 Hz stimulation chambers compared to habituation day in the ChR2 groups (n = 10-15, *p* < 0.01). ChR2 groups also preferred the stimulation chamber relative to YFP animals on the stimulation day (n = 10-12, *p* < 0.01). In contrast, female rats demonstrated no preference or aversion for the chamber paired with IL stimulation (n = 11-13/group, repeated-measures 2-way ANOVA: stimulation F_(1, 22)_ = 5.79, *p* < 0.05, stimulation x ChR2 F_(1, 22)_ = 0.005, *p =* 0.94). Additionally, IL stimulation did not affect general locomotor activity in either sex (**Fig. 2C**). Together, these findings indicate that the activity of male IL glutamatergic neurons has a positive affective valance as animals were motivated to seek out the stimulation. However, this cell group does not appear to modify affective state in females.

**Figure 2:**
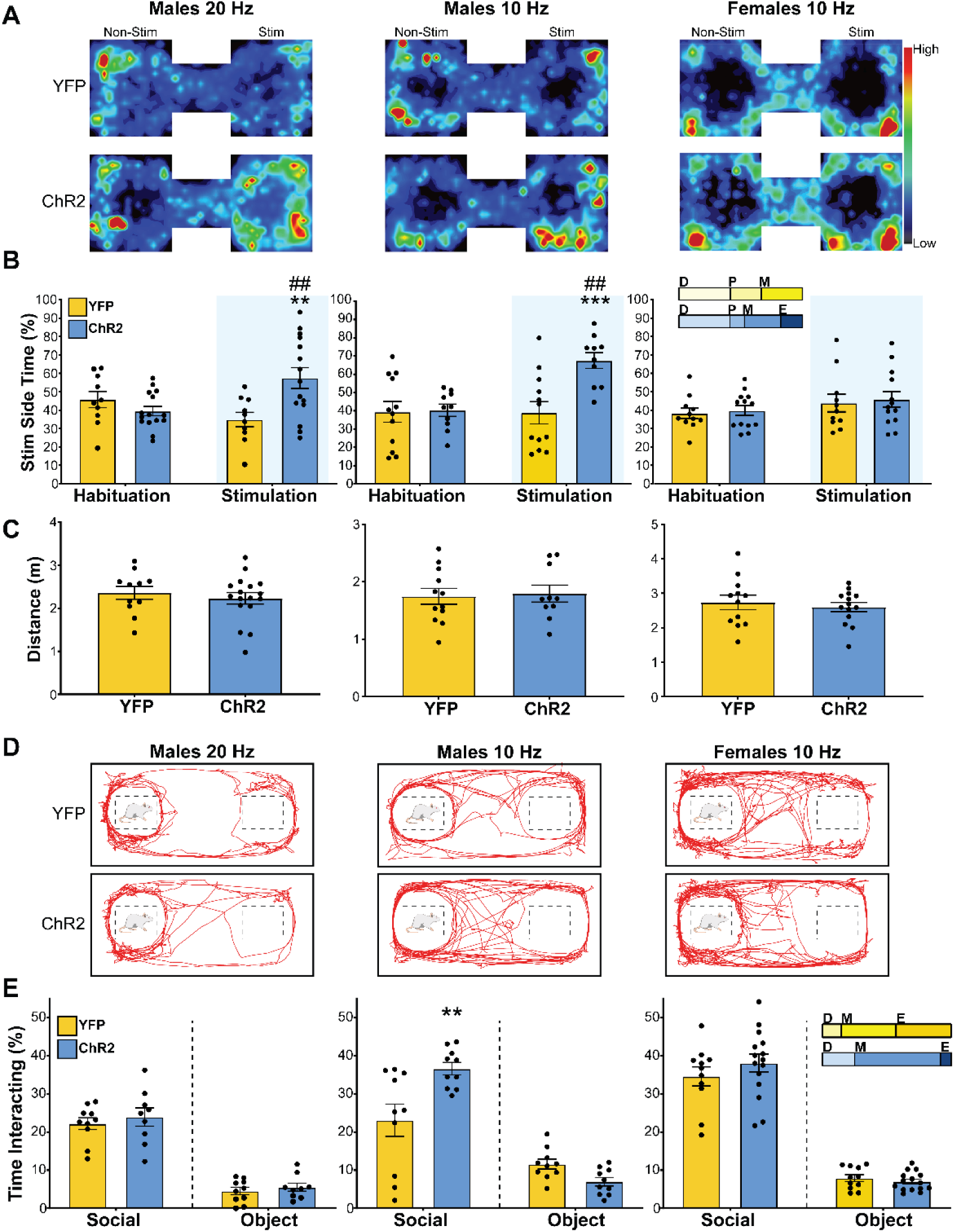
IL pyramidal neuron activity was preferred and increased social motivation in males but not females. **A**) Heat maps illustrate mean animal position in RTPP arena on stimulation day (n = 10-15/group). (**B**) Male ChR2 rats preferred the chamber paired with 20 Hz or 10 Hz stimulation (blue shading) relative to habituation day and YFP controls. Females showed no preference or aversion for IL stimulation. ** p < 0.01, *** p < 0.001 vs. YFP within stimulation. ^##^ p < 0.01 vs. habituation within ChR2. Inset: portion of rats in each estrous cycle phase. D: diestrus, P: proestrus, M: metestrus, E: estrous. (**C**) Total distance traveled during stimulation found no treatment-based differences in locomotion. (**D**) Representative movement traces during the social interaction test (n = 7-16/group). (**E**) Interaction, defined as nose-poke onto social cage, was increased by 10 Hz stimulation in males but not 20 Hz in males or 10 Hz in females. ** p < 0.01 vs. YFP. Inset: portion of rats in each estrous cycle phase.

### 3.3 Social behavior

The three-chamber social interaction assay was used to determine the influence of IL glutamatergic neurons on sociability. During the social motivation test (**Fig. 2D**), there was no difference in social interactions with 20 Hz stimulation in males (n = 9-10/group, unpaired t-test: ChR2 vs YFP t_(17)_ = 0.77, *p =* 0.45). However, 10 Hz stimulation in males increased social interactions (**Fig. 2E**; n = 10-12/group, unpaired t-test: ChR2 vs YFP t_(18)_ = 3.00, *p <* 0.01). In contrast, 10 Hz stimulation in females did not impact social interaction (n = 12-16/group, unpaired t-tests: ChR2 vs YFP t_(24)_ = 1.01, *p =* 0.32). Furthermore, neither male nor female IL stimulation affected preference for novel vs. familiar interactors (**Table S1**). Overall, these results indicate that IL glutamatergic activity increases male social motivation in a frequency-dependent manner; however, this cell population does not alter female social motivation.

### 3.4 Endocrine reactivity

To determine the effect of stimulating IL pyramidal neurons on neuroendocrine responses, blood glucose and plasma corticosterone were monitored during restraint stress. In males, optic stimulation decreased corticosterone (n = 10-17/group, mixed-effects: time F_(4,81)_ = 139.4, p < 0.0001, ChR2 F_(1,25)_ = 8.93, *p* < 0.01) at the 30-minute timepoint (**Fig. 3A**; *p* < 0.01). Additionally, corticosterone was decreased post-stimulation in the ChR2 group during stress recovery (90 min; *p* < 0.05). Stimulation also decreased blood glucose (mixed-effects: time F_(4,90)_ = 62.86, *p* < 0.0001, time x ChR2 *F*_(4,90)_ = 3.43, *p* < 0.05) in male rats during restraint (**Fig. 3B**; 15 min, *p* < 0.01; 30 min, *p* < 0.05). In contrast, stimulation did not alter plasma corticosterone in female rats (**Fig. 3C**; n = 10-17/group, mixed-effects: time F_(4,93)_ = 62.94, *p* < 0.0001) with effects of time limited to within treatment. Additionally, stimulation increased glucose responses to stress in female rats (n = 10-17/group, mixed effects: time F_(4,90)_ = 24.96, *p* < 0.0001) specifically at the 15-min timepoint (**Fig. 3D**; *p* < 0.05). Collectively, these results suggest that IL activity in males reduces HPA axis activation as well as glucose mobilization. In females, IL activity does not appear to alter HPA axis response to stress. However, IL stimulation in females increases glucose, possibly through sympathetic mobilization of epinephrine and/or glucagon.

**Figure 3:**
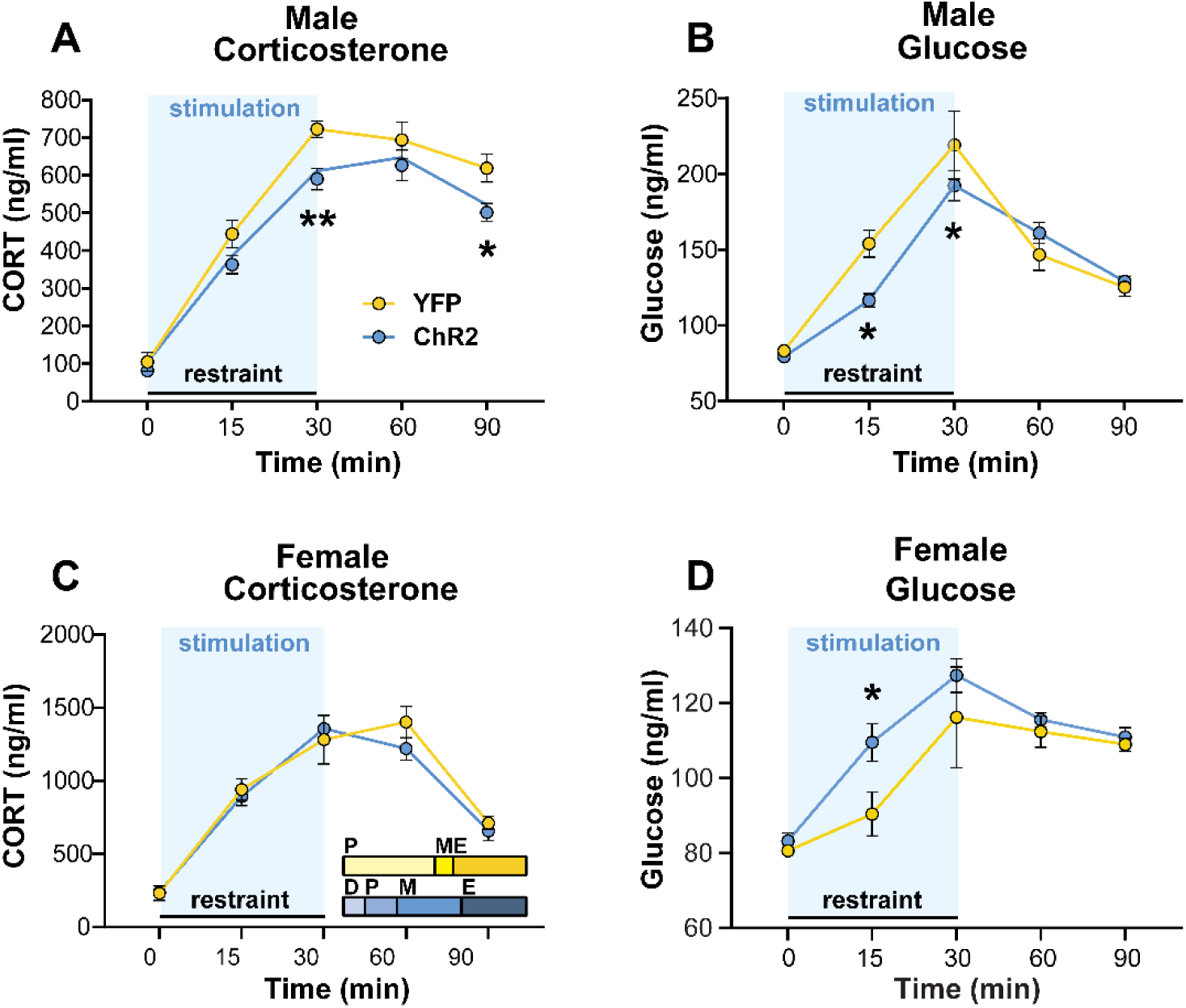
IL activation during acute restraint stress attenuated endocrine responses in males but increased female stress reactivity. Stimulation (blue shading) during restraint lowered plasma corticosterone **(A)** and blood glucose **(B)** in ChR2 males. **(C)** Stimulation in females did not alter corticosterone responses. Inset: portion of rats in each estrous cycle phase. D: diestrus, P: proestrus, M: metestrus, E: estrous. **(D)** IL stimulation during restraint increased glucose mobilization in ChR2 females. n = 10-17/group, * p < 0.05, ** p < 0.01 vs. YFP.

### 3.5 Cardiovascular reactivity

Baseline hemodynamic measures were recorded in the homecage prior to handling (**Fig. S4**). Animals were then connected to fiber optic patch cords and exposed to a brightly lit novel environment as a psychogenic stimulus to examine IL glutamatergic effects on cardiovascular reactivity. Over the course of the stressor, there were no differences in activity between male YFP (n = 7) and ChR2 (n = 8) rats (**Fig. 4A**; Mann-Whitney: U = 407, *p* = 0.53). In terms of hemodynamic responses, stimulation decreased HR reactivity in male ChR2 rats (mixed-effects: time F_(30,280)_ = 19.61, *p* < 0.0001, time x ChR2 F(_30,280_) = 2.06, *p* < 0.01), an interaction present at the 8-minute timepoint (**Fig. 4B**; *p* < 0.05). After the first minute of stimulation, arterial pressures including mean (mixed-effects: time F_(30,172)_ = 25.99, *p* < 0.0001, time x ChR2 F(30,172) = 1.86, *p* < 0.01), systolic (mixed-effects: time F_(30,213)_ = 27.61, *p* < 0.0001), and diastolic (mixed-effects: time F_(30,189)_ = 20.56, *p* < 0.0001, ChR2 F_(1,12)_ = 8.05, *p* < 0.05, time x ChR2 F_(30,189)_ = 2.38, *p* < 0.01) were decreased in male ChR2 rats. Post-hoc analysis indicated these effects were present across numerous timepoints (**Fig. 4C-E**; 7-28 minutes; *p* < 0.05). In contrast, stimulation increased MAP and DAP in minute 1 (*p* < 0.05). For females, there was no effect of ChR2 on activity in the novel environment (**Fig. 4F**; n = 7-8, Mann-Whitney: ChR2 vs YFP U = 352, *p* = 0.15). However, IL activation increased female HR responses (mixed effects: time F_(30,390)_ = 25.83, *p* < 0.0001) early in the stressor (**Fig. 4G**; 1-9 min; *p* < 0.05) as well as at 25 min (*p* < 0.05). Mixed effects analysis of arterial pressure found main effects of time for MAP (**Fig. 4H**; time F_(30,390)_ = 31.96, *p* < 0.0001) and SAP (**Fig. 4I**; time F_(30,390)_ = 35.01, *p* < 0.0001) without virus-specific effects. Taken together, these results indicate activation of male IL glutamate neurons decreases HR and arterial pressure responses to psychological stress. In contrast, activation in females increases HR.

**Figure 4:**
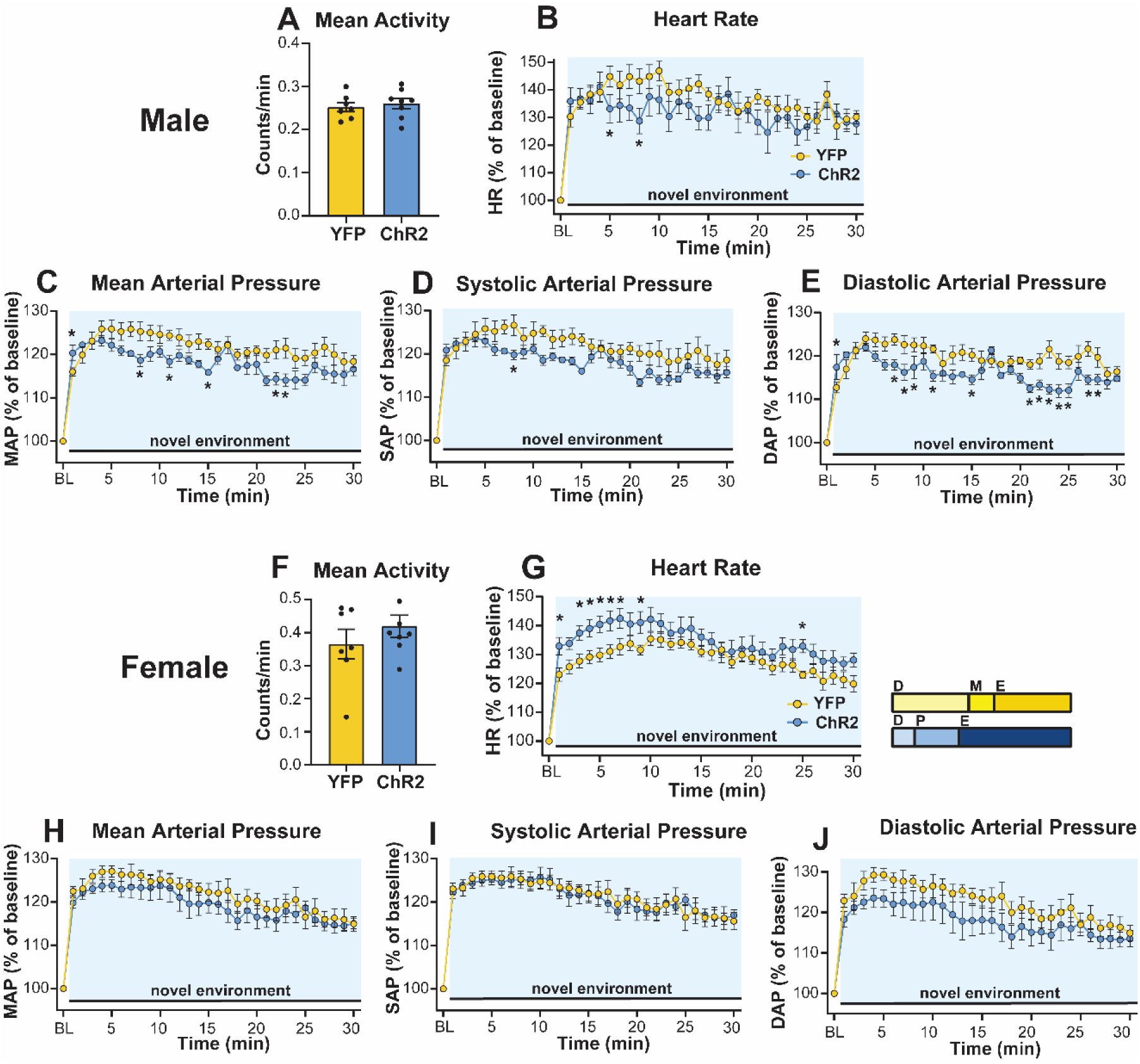
Activation of glutamatergic IL neurons during novel environment exposure reduced male cardiovascular stress responses but increased female HR reactivity. **(A)** Male IL activation did not affect mean activity in the novel environment. Despite an increase in MAP and DAP during the first minute of stimulation (blue shading) and stress, all recorded hemodynamic responses were decreased in ChR2 males **(B-E)**. **(F)** Stimulation did not alter activity in females. **(G)** Female IL stimulation elevated HR reactivity compared to YFP controls but did not affect arterial pressures **(H-J)**. Inset: portion of rats in each estrous cycle phase. D: diestrus, P: proestrus, M: metestrus, E: estrous. n = 7-8/group, * p < 0.05 vs. YFP.

### 3.6 Effects of chronic stress on cardiac function

Resting homecage LF/HF of HRV was recorded via radiotelemetry to assess how prior optic stimulation affected circadian autonomic balance and net cardiac sympathetic drive during CVS. In the absence of ongoing stimulation, ChR2 (n = 8) males had lower LF/HF than YFP (n = 7) rats (**Fig. 5A**; mixed-effects: time F_(5,62)_ = 4.45, *p* < 0.01, ChR2 F_(1,13)_ = 5.33, *p* < 0.05). In the AM the day prior to CVS, ChR2 males had lower baseline LF/HF (*p* < 0.05) leading to decreased sympathetic drive the morning of CVS day 14 (*p* < 0.05). In females, there were circadian effects on LF/HF (**Fig. 5B**; n = 5-6/group, mixed-effects: time F_(5,45)_ = 5.98, p < 0.01) but no differences due to virus (mixed-effects: ChR2 F_(1,9)_ = 1.28, p > 0.05).

**Figure 5:**
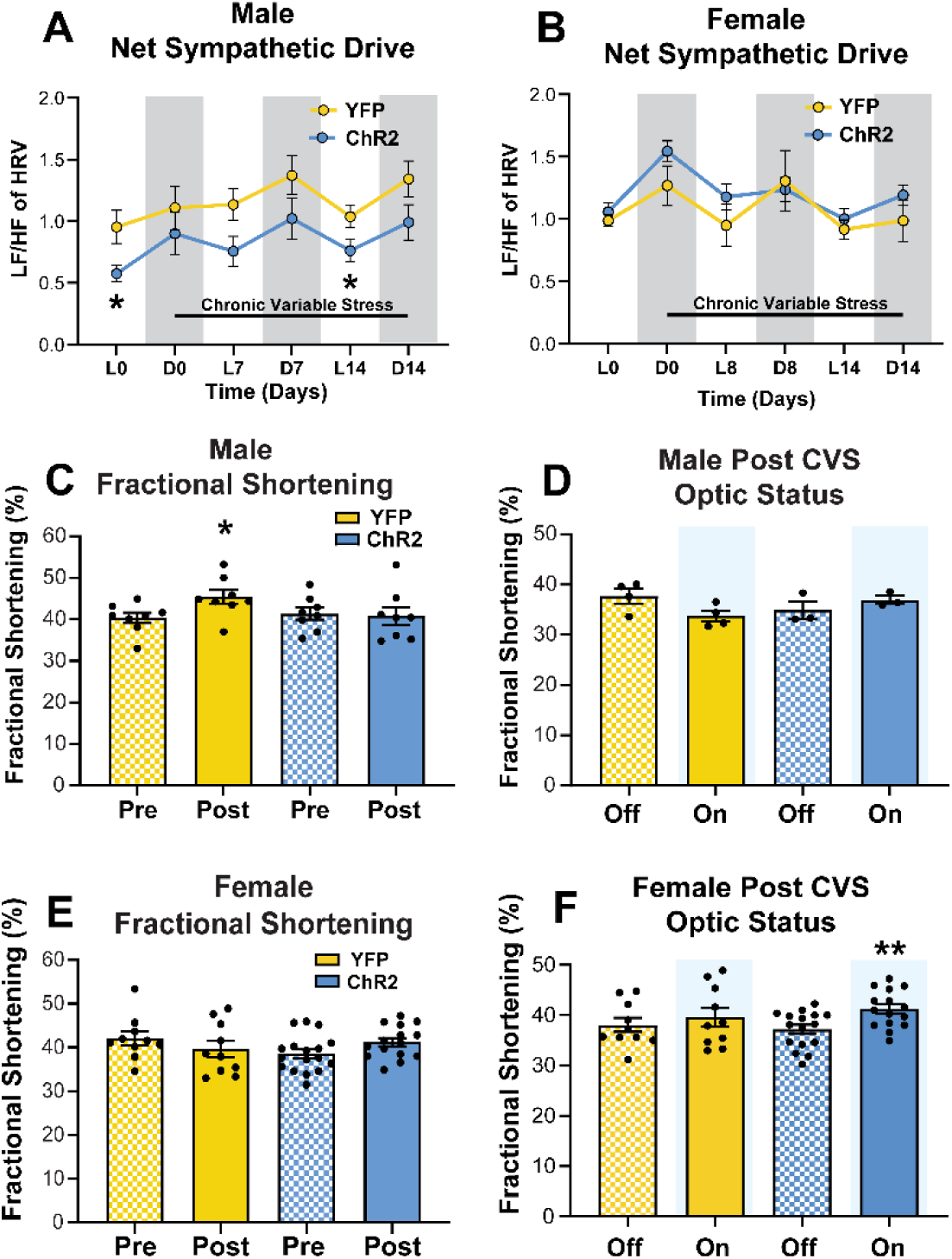
The effects of chronic stress and IL activity on cardiac function. **(A)** The ratio of LF to HF components of HRV were reduced in ChR2 males at baseline and the end of CVS. **(B)** Females had no treatment-based effects on net sympathetic drive. L: light phase, D: dark phase, LF: low frequency, HF: high frequency, HRV: heart rate variability. **(C)** CVS increased FS in YFP but not ChR2 males. **(D)** Acute optic status (On vs. Off) did not affect FS in males. **(E)** Females showed no changes in FS after CVS. **(F)** However, acute optogenetic activation (blue shading) increased female FS after CVS. * p < 0.05 vs YFP or Pre CVS YFP, ** p < 0.01 vs. ChR2 Off.

Left ventricle echocardiography was performed before and after CVS to examine the *in vivo* cardiac consequences of chronic stress and the effects of IL stimulation on cardiac contractility. Fractional shortening (FS) was measured as the portion of diastolic dimension lost in systole. In male rats treated with YFP (n = 8), CVS increased FS (**Fig. 5C**; Wilcoxon: Post vs Pre W = 30.0, *p* < 0.05). In contrast, there was no effect of CVS on FS in ChR2 (n = 8) males (Wilcoxon: Post vs Pre W = −4.0, *p* = 0.84). To determine whether post-CVS FS was affected by acute stimulation, measurements were taken with optics off and then on (**Fig. 5D**). Here, a cohort of males showed no effect of acute optic status on FS (n = 3-4/group, Wilcoxon: YFP On vs Off W = 4.0, *p* = 0.13; ChR2 On vs Off W = 2.0, *p* = 0.75). In females there was no effect of CVS on FS (**Fig. 5E**; n = 10, YFP Post vs Pre W = −21.0, *p* = 0.25; n = 17, ChR2 Post vs Pre W = 53.0, *p* = 0.07). However, optic stimulation increased post-CVS FS in ChR2 animals (**Fig. 5F**; Wilcoxon: On vs Off W = 87.0, *p* < 0.01). In aggregate, these findings indicate that a history of IL stimulation limits net cardiac sympathetic drive during chronic stress in males, an effect that may protect against CVS-increased FS. Conversely, neither CVS nor prior IL stimulation affected autonomic balance or cardiac function in females. However, acute IL stimulation after CVS increased female cardiac contractility.

### 3.7 Chronic stress effects on cardiac structure

Left ventricular morphological analysis was carried out to examine potential structural contributions to altered function. In YFP males (**Fig. 6A**; n = 8), CVS increased wall thickness of the posterior wall in systole (**Fig 6B**; Wilcoxon: Post vs Pre W = 28, *p* < 0.05) and the anterior wall in both systole (Wilcoxon: Post vs Pre W = 36, *p* < 0.01) and diastole (Wilcoxon: Post vs Pre W = 28, *p* < 0.05). Furthermore, increased wall thickness was sufficient to decrease ventricle size in systole (**Fig. 6C**; Wilcoxon: Post vs Pre W = −30, *p* < 0.05). Critically, male rats that had previously received IL stimulation (ChR2, n = 8) were protected from the effects of CVS on cardiac hypertrophy (**Fig 6D-E**; Wilcoxon: Post vs Pre W = −9 to 20, *p* > 0.05). In contrast to males, female rats (**Fig. 6F**) showed minimal effects of CVS on cardiac remodeling as YFP rats (n = 10) had decreased posterior wall thickness in diastole (**Fig. 6G**; Wilcoxon: Post vs Pre W = −37.0, *p* < 0.05) with no other structural changes (**Fig. 6H**; *p* > 0.05). Additionally, there were no significant changes in cardiac structure in the female ChR2 group (**Fig. 6I-J**; n = 16, Wilcoxon: Post vs Pre W = −45 to 25, *p* > 0.05). Overall, CVS-induced inward hypertrophic remodeling was prevented by prior IL activation in males. In contrast, female rats were generally resistant to the hypertrophic effects of chronic stress.

**Figure 6:**
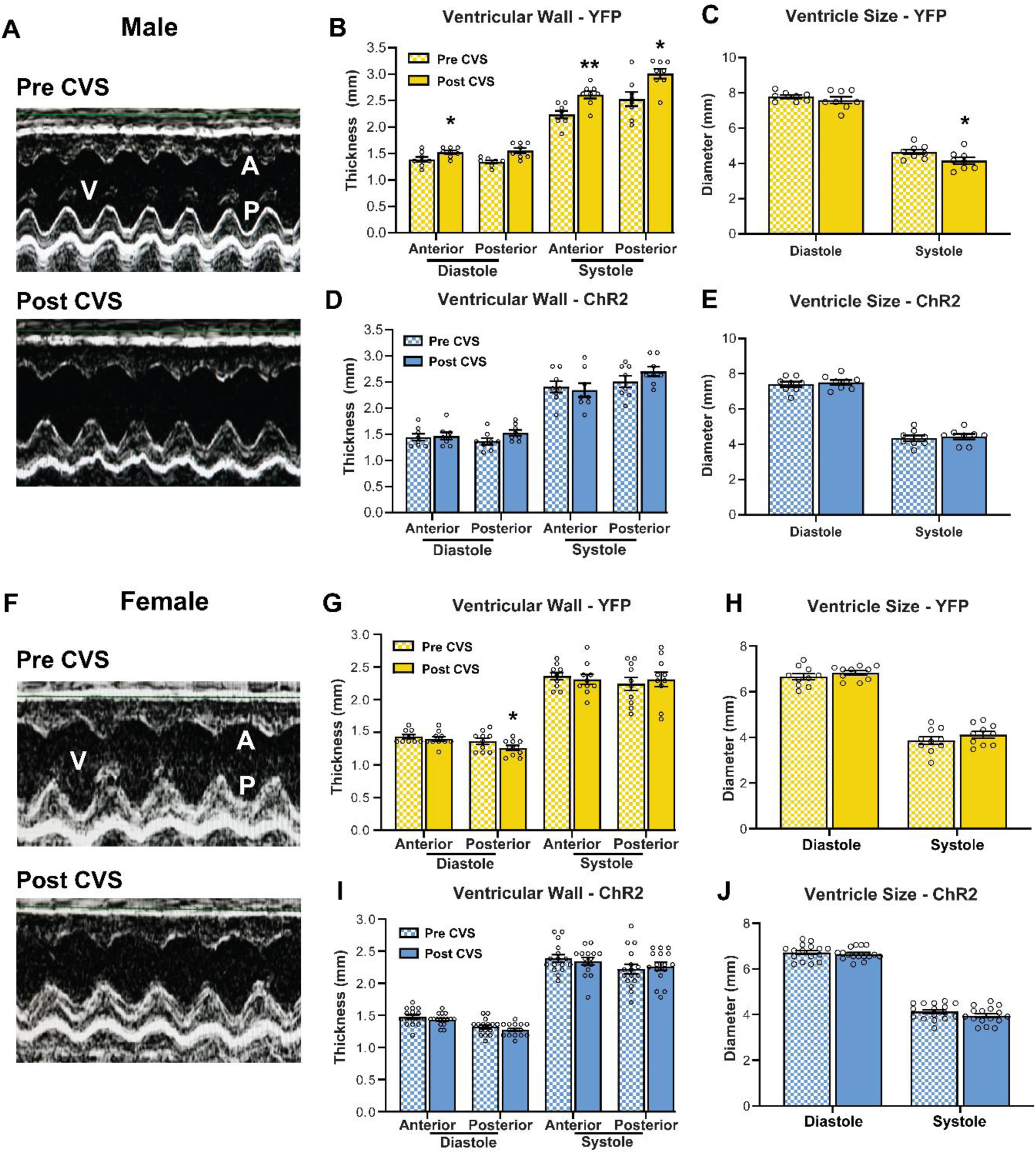
Effects of chronic stress and prior IL stimulation on cardiac structure. **(A)** Representative echocardiographic images of the left ventricle in bisected m-mode view before and after CVS in YFP males. V: ventricle, A: anterior wall, P: posterior wall. **(B)** CVS increased wall thickness in diastole and systole in YFP males, **(C)** reducing ventricle size. **(D, E)** ChR2 males that received prior stimulation were protected from the cardiac consequences of CVS. **(F)** Representative m-mode view of the left ventricle in YFP females before and after CVS. **(G, H)** Females did not exhibit ventricular hypertrophy after CVS as posterior wall thickness in diastole was decreased. **(I, J)** No myocardial structural changes were evident in ChR2 females. n = 8-16/group, * p < 0.05, ** p < 0.01 vs. Pre CVS YFP

## 4. Discussion

In the current study, optogenetic stimulation of glutamatergic IL pyramidal neurons was combined with behavioral, endocrine, and cardiovascular assessments. Our results show that, in males, IL pyramidal neuron activity was preferred, increased social motivation, and reduced acute physiological stress reactivity. Intriguingly, prior IL activation lowered net cardiac sympathetic drive and protected against subsequent myocardial remodeling after chronic stress. However, IL activity had fundamentally different regulatory effects in females. Stimulation did not have motivational valence or alter social behavior but increased acute physiological stress reactivity and cardiac contractility following chronic stress. Collectively, these findings identify sexual divergence in the cortical integration of affective and physiological systems, suggesting that vmPFC output signaling may differentially impact health outcomes in males and females.

The comorbidity of CVD and MDD shows sexual divergence with females at twice the risk (Goldstein et al., 2019; Möller-Leimkühler, 2007; Naqvi et al., 2005). Given the interactions between stress, mood disorders, and CVD, stress-reactive neural populations are well positioned to regulate affective and cardiovascular outcomes. Importantly, chronic stress exposure in male rats reduces IL pyramidal neuron dendritic arborization and increases local GABAergic signaling, suggesting reduced glutamatergic output (McKlveen et al., 2019, 2016; Radley et al., 2008). Further, long-term reduction of male IL glutamatergic output increases HPA axis activity and impairs vascular function (Myers et al., 2017; Schaeuble et al., 2019). Collectively, IL output neurons represent a target for modulating behavioral and physiological responses to stress. Thus, we sought to test this hypothesis through real-time *in vivo* assessments of behavior, physiology, and organ function. Altogether, we found that the neurobiology of emotional behaviors, endocrine-autonomic integration, and cardiovascular outcomes differed substantially between sexes.

The RTPP test was used to assess the affective state of experimental animals by measuring preference or aversion for IL neural activity. Here, male rats preferred the chamber paired with either 10 Hz or 20 Hz stimulation, but the valence of female IL activation was neither positive nor negative. Previous work indicates that male mPFC stimulation reduces social avoidance after social defeat (Covington et al., 2010), yet relatively few studies have examined IL function in females. Although social interaction induces less IL immediate early-gene expression in females than males (Mikosz et al., 2015; Stack et al., 2010), the current study is, to our knowledge, the first examination of IL regulation of female social behavior. IL activity did not alter social behavior in females, indicating a significant sexual divergence in the neural regulation of sociability. However, males exhibited frequency-dependent increases in social motivation. These results suggest that stimulation near the intrinsic pyramidal neuron firing rate (4-10 Hz) (Homayoun and Moghaddam, 2007; Ji and Neugebauer, 2012) is required to increase male social motivation.

Activation of IL glutamatergic neurons also reduced male glucose and corticosterone responses to stress. Conversely, female IL glutamatergic stimulation increased glucose mobilization without affecting corticosterone, suggesting a role in sympatho-excitation. Female IL activity also increased tachycardic responses to stress, while male IL stimulation reduced cardiovascular stress responses including HR and arterial pressures. These experiments indicate that IL activity has opposing effects on endocrine-autonomic integration in males and females whereby male glutamatergic IL neurons cause widespread inhibition of the stress response and female IL neurons facilitate sympathetic responses.

Chronic stress exposure induced ventricular hypertrophy and increased endocardial FS in males, suggesting that increased wall thickness likely accounted for the increase in FS. This effect was prevented by a history of IL stimulation in males, likely arising from a reduction in symaptho-vagal imbalance throughout CVS. In fact, previous IL stimulation limited resting net sympathetic drive, a major risk for CVD (Thayer et al., 2012; Wulsin et al., 2015). Given that rats received no optic stimulation during CVS and that acute optic status did not impact male FS, this effect was likely driven by stimulation-induced IL plasticity. IL stimulation has been shown to induce persistent morphological changes in males including increased excitatory synapses onto pyramidal neurons, suggesting a prolonged state of enhanced excitability (Fuchikami et al., 2015; Moda-Sava et al., 2019). Thus, lower resting net sympathetic tone and/or reduced sympathetic activity during stressors may have prevented the consequences of chronic stress to induce inward hypertrophic remodeling of the myocardium. These effects were not present in females as hypertrophic remodeling was not evident and acute stimulation increased FS. Collectively, these results indicate that prior male IL glutamatergic activity is sufficient to restrain responses to chronic stress while female IL activity increases cardiac contractility independent of structural changes.

Physiological responses to stress are critical for survival, tightly defended, and limited in maximum capacity; accordingly, our manipulations only caused moderate changes in response magnitude to acute challenges. However, our findings suggest that the interaction of altered stress responding with chronic stressors is sufficient to account for differences in cardiac structure and function. It is also worth noting that prior studies on the role of vmPFC in stress responding have yielded equivocal results. Specifically, lesions studies in male rats have observed both increased (Diorio et al., 1993; Figueiredo et al., 2003) and decreased (Radley et al., 2006) neuroendocrine stress responding. Similarly, pharmacological studies found that male vmPFC both reduces (Müller-Ribeiro et al., 2012) and enhances (Tavares et al., 2009) cardiovascular stress responses. These discrepancies are not limited to rodents as pharmacological studies in non-human primates suggest that vmPFC increases cardiovascular stress responding (Alexander et al., 2020), while electrical stimulation in humans causes hypotension (Lacuey et al., 2018). Although it is possible that variations in acute stressors, selectivity of cortical subregion targeting, and/or species may account for these divergent findings, all of the described studies used non-specific blockade or stimulation. Given the complexity of cortical circuitry and the numerous local inhibitory mechanisms (McKlveen et al., 2019), cell-type specific targeting has the ability to isolate the precise contribution of vmPFC principal cells independent of afferents, interneurons, and glia. To address this issue, our prior studies developed anatomical and cell-type specific approaches to reduce glutamate outflow from IL pyramidal neurons. Genetic knockdown of IL glutamate release in males increased neuroendocrine and cardiovascular responses to both acute and chronic stress (Myers et al., 2017; Schaeuble et al., 2019), demonstrating the necessity of IL output for inhibiting physiological stress responses. The current cell-type specific optogenetic targeting adds further temporal specificity and identifies the sufficiency of male IL glutamate neurons for stress inhibition. Albeit with opposing effects in females.

The IL does not directly innervate the neurons that govern endocrine and autonomic stress responses. Accordingly, downstream glutamate signaling from IL synaptic terminals requires intermediary synapses. The exact circuits engaged by IL pyramidal cells to bring about the observed effects remain to be determined. However, anterograde mapping studies indicate that IL projections widely innervate the forebrain and brainstem (Vertes, 2004; Wood et al., 2018). We previously found that, in males, stress-activated IL neurons innervate local inhibitory GABAergic neurons in the posterior hypothalamus (PH) (Myers et al., 2016). Furthermore, blocking GABAergic tone in the PH reduces social behavior and increases HPA axis reactivity suggesting inhibition of the PH may be important for limiting behavioral and physiological stress responses (Myers et al., 2016). Additionally, IL inputs to the amygdala are critical for fear extinction and reducing anxiety-like behavior (Adhikari et al., 2015; Sierra-Mercado et al., 2011). Interestingly, amygdala-projecting IL neurons are both resistant to stress-induced dendritic retraction as well as sensitive to estrogen (Shansky et al., 2010, 2009). Thus, the IL-amygdala circuit could play a key role in sex differences in behavioral regulation. Although the downstream mechanisms of IL cardiovascular regulation are unknown, male IL projections target pre-autonomic cell groups in the brainstem and give rise to multi-synaptic pathways that innervate the adrenal medulla (Dum et al., 2019; Gabbott et al., 2005). Further sex-specific analysis of IL synaptic signaling in forebrain and brainstem nuclei is necessary to determine the basis of divergent behavioral and physiological integration.

While ovarian hormones have far-reaching effects on behavior and physiological systems, we did not control for estrous cycle phase. Instead, we used randomly cycling females. Cycle phase was reported for each treatment, but statistical power was insufficient to examine phase as a factor. It remains to be determined how gonadal hormones might contribute to the sexually divergent effects observed. Estrogen receptor (ER) α, β, and the g-protein coupled ER are expressed in pyramidal and non-pyramidal PFC neurons in both sexes (Almey et al., 2014; Montague et al., 2008). Further, ER localized to PFC dendritic spines regulates synaptic morphology and ionotropic glutamate receptor ubiquitination/degradation following repeated stress (Hao et al., 2006; Wei et al., 2014; Yuen et al., 2016). These protective effects are dependent on PFC ERα and estradiol-synthesizing aromatase (Wei et al., 2014), indicating a role for extra-ovarian estrogen synthesis. In addition, ER expressed in axons and axonal terminals rapidly alters pre-synaptic transmission in pyramidal cells (Almey et al., 2014). Although studies have described multiple interactions between ER and cortical glutamate signaling (Galvin and Ninan, 2014; Hara et al., 2018), much less is known about PFC progesterone signaling. Progesterone receptors are expressed in frontal cortex (Guerra-Araiza et al., 2003) but progesterone derivatives also signal through GABA_A_ receptors and regulate GABA subunit expression (Andrade et al., 2012). Moreover, sex differences in IL regulation of behavior and physiology could arise from the actions of androgens. Androgen receptors (AR) are expressed in PFC neurons and glia (Finley and Kritzer, 1999). AR is also enriched in VTA-projecting PFC neurons that influence extracellular dopamine in PFC through downstream VTA glutamate signaling (Aubele and Kritzer, 2012). It is particularly interesting that androgens and estrogens act in opposition to modify PFC dopamine, norepinephrine, and serotonin metabolism during novel environment stress (Handa et al., 1997). Thus, gonadal steroids and locally synthesized modulators affect PFC cellular processes and projection activity, ultimately engaging multiple neurotransmitter systems. Accordingly, the specific cellular and synaptic processes contributing to sex-dependent stress reactivity and resilience are promising avenues for identifying therapeutic targets.

The current study identified a cortical node integrating mood-related behaviors with cardiovascular outcomes. Moreover, activity in this vmPFC cell population produced sex-specific effects on behavior and physiology. In addition to highlighting the necessity of sex-based investigation, these data point to a neurochemical basis for sex differences in stress-related health determinants. Ultimately, further investigation of brain-body interactions in the face of prolonged stress may provide a better understating of disease risk and resilience factors.

## Funding

This work was supported by NIH grants R00 HL122454 and R01 HL150559 to Brent Myers. Data generated by the University of Cincinnati Mouse Metabolic Phenotyping Core were supported by NIH grant U2C DK059630.

## Author contributions

**Tyler Wallace:** Formal analysis, Investigation, Validation, Visualization, Writing - original draft, Writing - review & editing. **Derek Schaeuble:** Formal analysis, Investigation, Validation, Visualization, Writing - original draft, Writing - review & editing. **Sebastian A. Pace:** Formal analysis, Investigation, Validation. **Morgan K. Schackmuth:** Formal analysis, Validation. **Shane T. Hentges:** Methodology, Resources, Supervision, Writing - review & editing. **Adam J. Chicco:** Formal analysis, Investigation, Methodology, Resources, Writing - review & editing. **Brent Myers:** Conceptualization, Funding acquisition, Investigation, Methodology, Resources, Supervision, Writing - original draft, Writing - review & editing.

## Disclosures

The authors have no conflicts of interest to disclose.

## Acknowledgments

AAV5 vectors were provided by the University of North Carolina Vector Core under material transfer agreement with Karl Deisseroth and Stanford University. This article was first published as a preprint: Wallace T, Schaeuble D, Pace SA, Schackmuth MK, Hentges ST, Chicco AJ, Myers B. Sexually divergent cortical control of affective-autonomic integration. *bioRrxiv* doi.org/10.1101/2020.09.29.319210

## Supplemental Data

### Injection and cannula placement

Viral vectors were injected in the IL. Fiber optic cannulas were targeted to the dorsal boundary of the IL, allowing cone-shaped illumination ventral to the fiber tip to stimulate IL CaMKIIα-positive neurons (**Fig. S1**).

**Figure S1:**
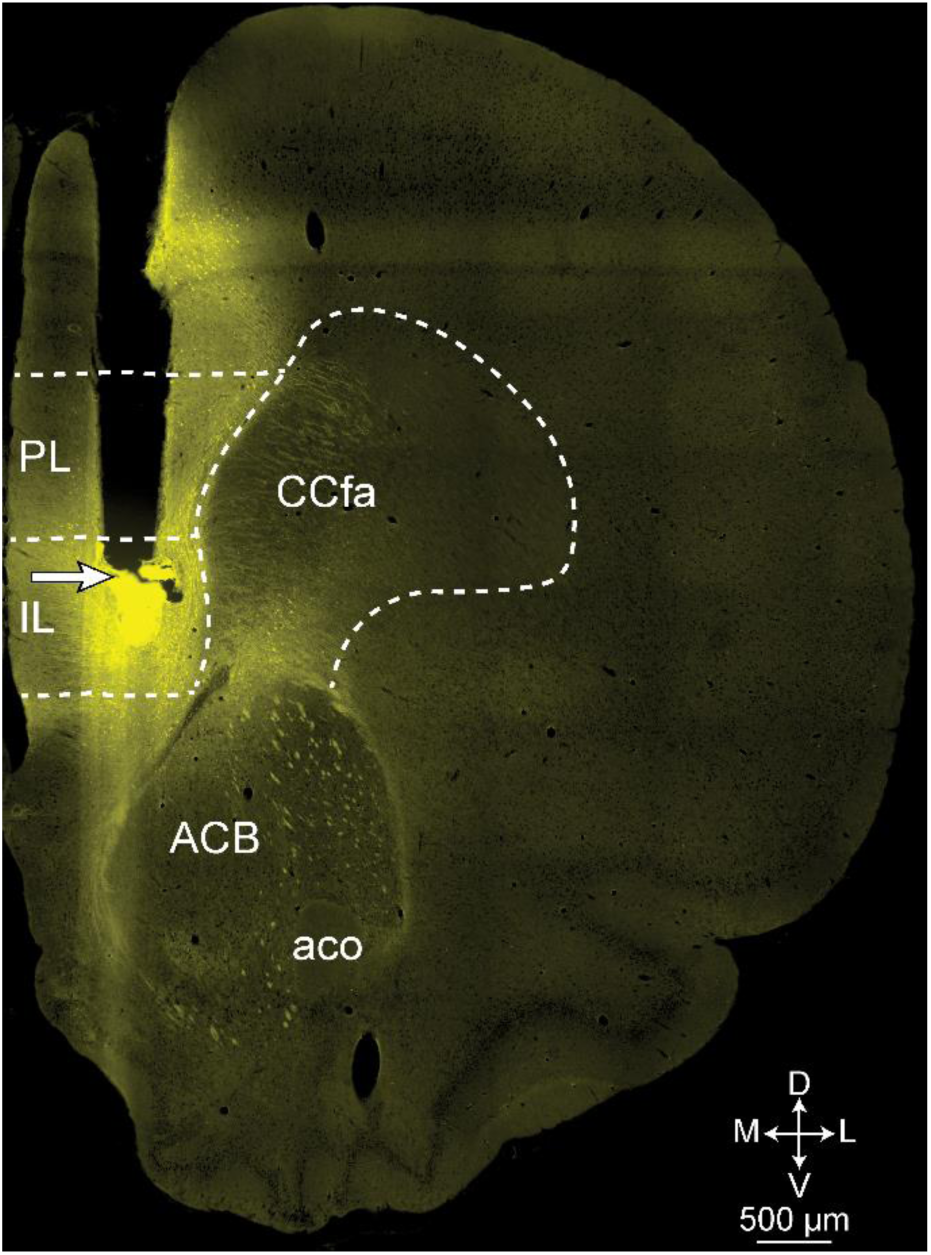
Photomicrograph of IL cannulation site. 10x image of cannula tract targeting the IL with AAV-delivered YFP fluorescence. White arrow indicates location of fiber tip. Dashed white lines indicate region borders. ACB - nucleus accumbens, aco – anterior commissure, CCfa – corpus callosum anterior forceps, IL – infralimbic, PL-prelimbic.

### Stimulation-induced c-Fos expression

To determine the impact of light stimulation on neural activity, we quantified c-Fos positive nuclei labeled through chromogen immunohistochemistry (**Fig. S2**). Stimulation increased c-Fos positive nuclei in males (n = 6-7/group, Welch’s unpaired t-test: ChR2 vs YFP t_(6.416)_ =3.268, *p <* 0.05) and females (n = 6-7/group, unpaired t-test: ChR2 vs YFP t_(11)_ = 4.475, *p <* 0.005). These results indicate, that at the termination of experimentation, 10 Hz, 1 mW pulsatile stimulation significantly increased cellular activation in the IL of both sexes. While the mean value of c-Fos positive cells appears higher in males, there was no significant (p = 0.08) sex difference between the ChR2 groups.

**Figure S2:**
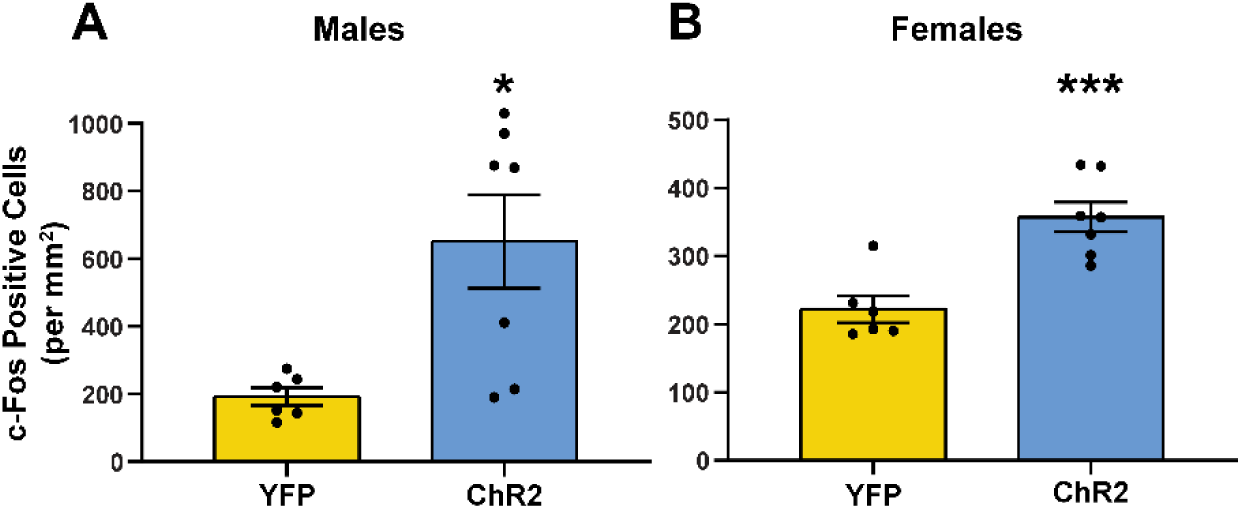
Optogenetic stimulation increased c-Fos positive cell count in males and females. At the conclusion of experiments 2 and 3, rats received 5 minutes of optic stimulation (10 Hz, 5 ms pulses, 1 mW) followed by 90 minutes of recovery prior to tissue collection. Chromogen immunohistochemistry was used to identify and quantify c-Fos positive cells in the IL. * p < 0.05, *** p < 0.005 vs YFP within sex.

### Social novelty preference

Time spent interacting with novel and familiar rats. Stimulation of IL pyramidal neurons did not affect preference for social novelty in males or females.

**Table S1:** Social novelty preference.

| Experiment | Virus | Novel interaction (% time) | Novel interaction (SEM) | Familiar interaction (% time) | Familiar interaction (SEM) | n |
| --- | --- | --- | --- | --- | --- | --- |
| **Male 20 Hz** | YFP | 16.63 | 1.27 | 13.06 | 2.81 | 7 |
|  | ChR2 | 12.84 | 1.72 | 9.73 | 0.94 | 10 |
| **Male 10 Hz** | YFP | 16.43 | 2.07 | 13.13 | 2.17 | 12 |
|  | ChR2 | 18.19 | 1.81 | 16.32 | 2.47 | 10 |
| **Female 10 Hz** | YFP | 24.84 | 2.61 | 16.56 | 1.62 | 12 |
|  | ChR2 | 21.58 | 1.55 | 16.96 | 1.51 | 16 |

### Homecage stimulation

Following behavioral experiments, male rats from experiment 2 and female rats from experiment 3 went through a homecage stimulation protocol the week prior to novel environment. Each rat was removed from the homecage, tethered to a fiber optic patch cord, and then returned to the cage. After 2 minutes, optics were turned on (0.5 mW, 5 ms pulses, 5 Hz) for one minute then turned off for one minute. This procedure repeated for 10 Hz and 20 Hz stimulation before power was increased to 1.3 mW and the protocol repeated. In total, each rat was tethered for 14 minutes and received stimulation for 6 minutes. For male rats, heart rate (HR) and mean arterial pressure (MAP) were recorded. Photostimulation of the IL did not affect HR or MAP (**Fig. S3**). Specifically, the 10 Hz stimulation used for cardiovascular reactivity studies had no effect at 0.5 or 1.3 mW on HR (n = 7-8, Mann-Whitney: ChR2 vs YFP 10 Hz 0.5 mW U = 25, *p* = 0.78; ChR2 vs YFP 10 Hz 1.3 mW U = 24, *p* = 0.69) or MAP (n = 7-8, Mann-Whitney: ChR2 vs YFP 10 Hz 0.5 mW U = 27, *p* = 0.96; ChR2 vs YFP 10 Hz 1.3 mW U = 21, *p* = 0.46). Although, there was an overall trend for decreased HR and MAP as a function of time post-handling. Hemodynamic parameters were not recorded in females.

**Figure S3:**
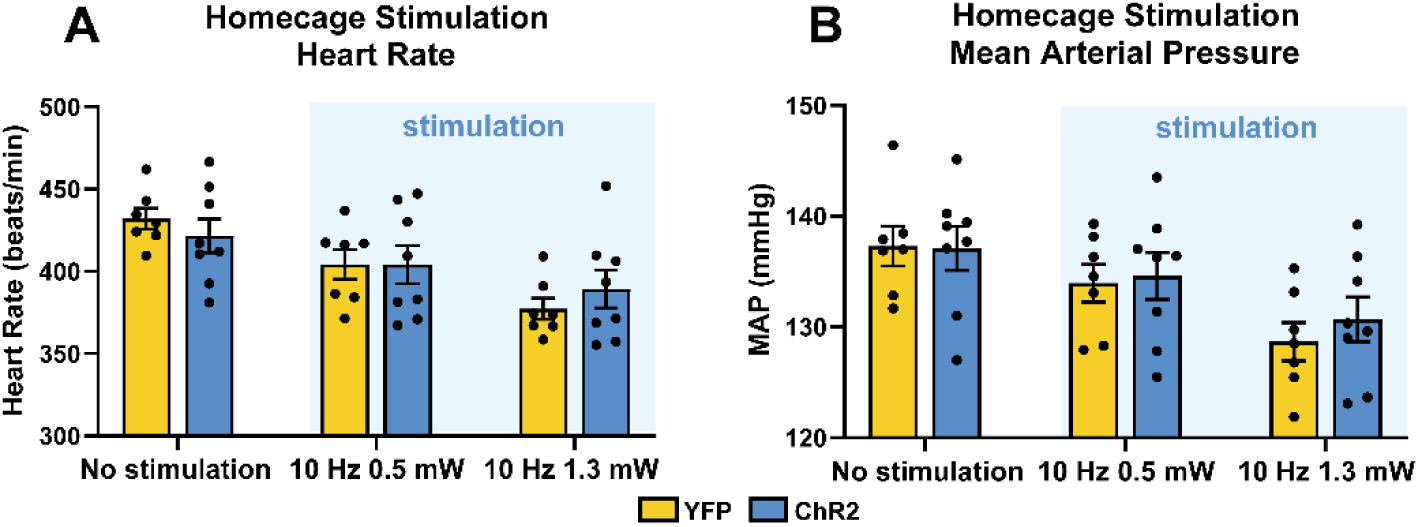
Optogenetic stimulation did not affect homecage HR or MAP in non-stressed male rats. HR and MAP were recorded over 2-minute periods of 10 Hz stimulation (0.5 and 1.3 mW).

### Baseline hemodynamic measures

The weekend prior to novel environment, baseline cardiovascular measurements were recorded at approximately the same time of day that experiments were conducted. Data were collected while rats were untethered in their homecages without optic stimulation (**Fig. S4**). In males, HR (n = 7-8, Mann-Whitney: ChR2 vs YFP U = 19, *p* = 0.34) and MAP (n = 7-8, Mann-Whitney: ChR2 vs YFP U = 23, *p* = 0.59) were not different. Likewise, female HR (n = 7-8, Mann-Whitney: ChR2 vs YFP U = 25, *p* = 0.78) and MAP (n = 7-8, Mann-Whitney: ChR2 vs YFP U = 28, *p* > 0.99) had no differences.

**Figure S4:**
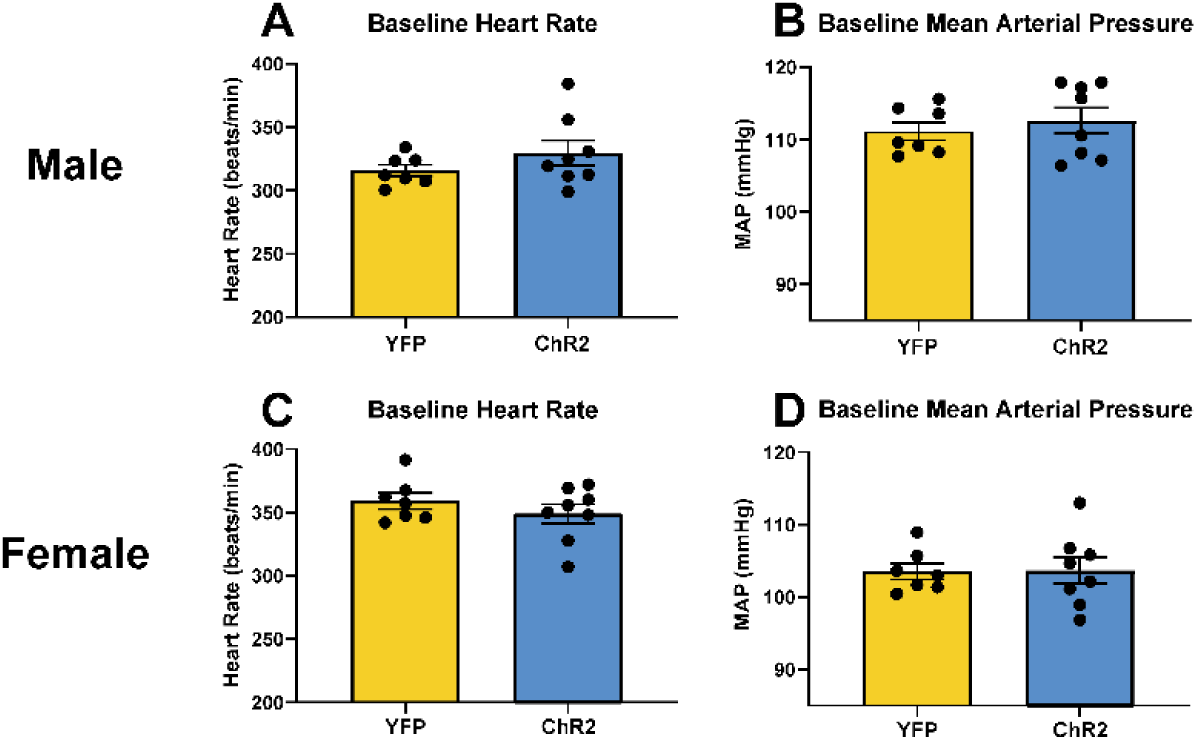
Baseline HR and MAP in male and female rats. Male rats **(A,B)** had no differences in baseline HR or MAP (p > 0.05). In female rats **(C,D)**, there were also no differences in HR or MAP (p > 0.05).

## Notes

### Competing Interest Statement

The authors have declared no competing interest.

